# Multilevel neural gradients reflect transdiagnostic effects of major psychiatric conditions on cortical morphology

**DOI:** 10.1101/2021.10.29.466434

**Authors:** Bo-yong Park, Valeria Kebets, Sara Larivière, Meike D. Hettwer, Casey Paquola, Daan van Rooij, Jan Buitelaar, Barbara Franke, Martine Hoogman, Lianne Schmaal, Dick J. Veltman, Odile van den Heuvel, Dan J. Stein, Ole A. Andreassen, Christopher R. K. Ching, Jessica Turner, Theo G. M. van Erp, Alan C. Evans, Alain Dagher, Sophia I. Thomopoulos, Paul M. Thompson, Sofie L. Valk, Matthias Kirschner, Boris C. Bernhardt

## Abstract

It is increasingly recognized that multiple psychiatric conditions are underpinned by shared neural pathways, affecting similar brain systems. Here, we assessed *i)* shared dimensions of alterations in cortical morphology across six major psychiatric conditions (autism spectrum disorder, attention deficit/hyperactivity disorder, major depression, obsessive-compulsive disorder, bipolar disorder, schizophrenia) and *ii)* carried out a multiscale neural contextualization, by cross-referencing shared anomalies against cortical myeloarchitecture and cytoarchitecture, as well as connectome and neurotransmitter organization. Pooling disease-related effects on MRI-based cortical thickness measures across six ENIGMA working groups, including a total of 28,546 participants (12,876 patients and 15,670 controls), we computed a shared disease dimension on cortical morphology using principal component analysis that described a sensory-fugal pattern with paralimbic regions showing the most consistent abnormalities across conditions. The shared disease dimension was closely related to cortical gradients of microstructure and intrinsic connectivity, as well as neurotransmitter systems, specifically serotonin and dopamine. Our findings embed the shared effects of major psychiatric conditions on brain structure in multiple scales of brain organization and may provide novel insights into neural mechanisms into transdiagnostic vulnerability.

## Introduction

Mental illness refers to a wide range of psychiatric conditions, affecting individuals, families, and health systems at large [1]. While conventional psychiatric nosology classifies mental illness into distinct categories mainly based on descriptive symptoms and behaviors [2], high co-occurrence of symptoms across disorders as well as transdiagnostic risk factors have prompted reconceptualization of mental illnesses along symptom dimensions [3–8]. The dimensional framework benefits detailed characterization of individual variations, and may allow for more direct brain-behavior associations than classic case-control comparisons that capture multiple symptom classes and mask clinical heterogeneity.

The shared components across major psychiatric diagnosis may be more clearly distinguishable at the neural level [4, 9], as the behavioral level likely involves complex interactions with society and the environment [10]. Structural magnetic resonance imaging (MRI), in particular, offers high spatial precision to help resolve the pattern of shared transdiagnostic effects across the cortical surface [4, 11–16]. A large body of prior case-control studies has reported reproducible patterns of structural alterations in cohorts with psychiatric diagnoses relative to controls [17–21], often pointing to widespread changes in cortical morphology in these conditions. More recently, efforts have been expanded to a transdiagnostic perspective, aiming to identify structural compromise that are shared across different diagnoses [22–24]. To ensure sensitivity of such efforts and to strengthen reproducibility, it becomes increasingly relevant to pool these investigations across multiple sites. One such initiative, spearheaded by the Enhancing NeuroImaging Genetics through Meta-Analysis (ENIGMA) consortium, has aggregated MRI and phenotypic data in thousands of healthy individuals and those with a psychiatric diagnosis [25]. Moreover, dedicated ENIGMA working groups have confirmed neuroanatomical disruptions in major psychiatric indications, including autism spectrum disorder (ASD) [26], attention deficit hyperactivity disorder (ADHD) [27], major depressive disorder (MDD) [28], obsessive-compulsive disorder (OCD) [29], bipolar disorder (BD) [30], and schizophrenia (SZ) [31], pointing to widespread changes in cortical morphology in each of these different conditions.

In addition to providing robust evidence of neuroanatomical signatures associated with each of these conditions, an emerging body of studies has pooled data across different indications to identify shared anomalies of psychiatric conditions [32, 33]. In an effort to identify factors contributing to the topography of cross-disorder brain changes, a recent study has taken this approach one step further and examined associations to *post mortem* gene expression data, searching for spatially co-varying gene lists that may carry susceptibility to transdiagnostic disease effects. This study identified that transdiagnostic effects may specifically be present in regions with greater expression of CA1 pyramidal genes that were suggested to play a role in regulating cortical thickness. Beyond these molecular risk factors, there is a broad range of cellular, metabolic, and functional properties of brain regions that may contribute to the regional susceptibility of transdiagnostic disease effects. An influential theory, also referred to as the structural model, posits that the internal microstructural and connectional markup of different brain regions, in particular their laminar differentiation and cortico-cortical connectivity patterns, may represent mesoscale features associated with the potential of a region to show plasticity, and to be susceptible to pathological processes [34]. According to this framework, paralimbic cortices with low laminar differentiation and higher-order connectivity profiles may be more susceptible to effects of neurological as well as psychiatric disorders. Here, we tested this approach, by aligning transdiagnostic effects with maps of microstructural variations derived from both *in vivo* imaging and 3D *post mortem* histology [35–38]. In recent work, the application of non-linear eigenvector decomposition to these datasets identified a “sensory-fugal” gradient that radiates from sensory and motor areas with strong laminar differentiation and higher myelination towards heteromodal association and paralimbic regions with less clear lamination and lower myelin content. Of note, similar gradients have also been derived from the analysis of intrinsic functional connectivity patterns obtained from resting-state functional MRI [37–39]. In line with foundational neuroanatomical conceptualization [34, 40, 41], an emerging literature has underscored a correspondence between such data-driven sensory-fugal gradients, and region-to-region variations in cortical plasticity and genetic control [39, 42–46], suggesting that these likely help understand susceptibility to disease as well [39, 42, 47–51].

The study of micro- and macroscale cortical organization as well as the identification of factors contributing to disease-related susceptibility for psychiatric conditions can be further complemented by studying associations to the neurotransmitter architecture of the human brain. Recent work based on *in vitro* receptor autoradiography in non-human primates has suggested that neurotransmitter systems are likely organized along similar gradients as cortical microstructure and connectivity, enabling on the one hand rapid and reliable information processing in sensory areas on the one hand, and slow, flexible integration of information in higher cognitive areas. Until similar resources become available in humans, one can approximate the spatial distributions of different neurotransmitter systems *in vivo*, based on the aggregation of positron emission tomography (PET) and single photon computed emission tomography (SPECT) studies sensitive to different receptor an transporter types [52–58]. Such mapping can thus provide a molecular perspective to complement microstructural and functional connectivity contextualization of transdiagnostic findings, promising new insights into factors contributing to the susceptibility of the brain to effects of different psychiatric conditions.

Here, we studied the association between multiscale neural organization and transdiagnostic effects on cortical morphology across six major psychiatric conditions, which represent a broad range of common and severe neurodevelopmental indications (ASD, ADHD, MDD, OCD, BD, and SZ). Aggregating data from thousands of patients and healthy controls previously studied across several ENIGMA working groups [26–31], we defined shared effects using principal component analysis, adapting a previous framework [32], and then associated the effects across multiple neural scales, namely (i) *in vivo* myeloarchitecture and intrinsic functional connectivity, (ii) *post mortem* 3D cytoarchitecture, and (iii) *in vivo* maps of neurotransmitter distributions.

## Results

### Study overview and participants

We obtained case-control maps of cortical thickness differences in patients relative to controls, resulting from several ENIGMA meta-analyses provided by a previous study, aggregating a total of 28,546 participants across 145 independent cohorts (1,821 ASD, 1,815 ADHD, 2,695 MDD, 2,274 OCD, 1,555 BD, 2,716 SZ; 15,670 site-matched controls **Table S1**) [32]. We then associated principal dimensions of morphological abnormalities with (i) *in vivo* myeloarchitecture and functional connectivity gradients obtained from the Human Connectome Project (HCP) [59], (ii) *post mortem* cytoarchitecture, by cross-referencing data to a ultra-high resolution 3D histological reconstruction of a human brain [60], and (iii) *in vivo* neurotransmitter topographies provided by PET/SPECT studies [52–58]. Approaches are openly available and replicable via the ENIGMA toolbox (https://enigma-toolbox.readthedocs.io) [61]. See *Methods* for more details.

### Shared dimensions of structural alterations across psychiatric conditions

Following standardized ENIGMA protocols (http://enigma.ini.usc.edu/protocols/imaging-protocols/), gray matter thickness for 68 cortical regions of the Desikan-Killany atlas [62] was calculated, and meta-analytic between-group differences in cortical thickness were assessed using inverse variance-weighted random-effects models (**Fig. 1A**) [32]. Using principal component analysis adopted in a recent study [32], we then estimated the shared disease dimensions explaining structural alterations across six conditions (**Fig. 1B**). The first dimension/component explained 55.7% of variance, and differentiated sensory/motor systems having positive scores from transmodal/paralimbic areas with negative scores (for *details*, and information on the other dimensions/components, see **Fig. S1A**). Stratifying the first dimension according to intrinsic functional communities [63], it indeed differentiated somatomotor/visual from default/frontoparietal/limbic networks (**Fig. 1B**). Similar spatial patterns were observed across the levels of the putative primate cortical hierarchy [40], differentiating idiotypic/unimodal from heteromodal/paralimbic levels. Notably, scores on the principal dimension translated into mean effect sizes across case-control analyses, with paralimbic regions showing strongest atrophy in patients relative to controls, while sensory/motor regions showed the least gray matter alterations (**Fig. S1B**). We also directly ran principal component analysis on previously reported effect size maps (Cohen’s d) concatenated across disorders, sourced from the ENIGMA toolbox [61] (**Fig. S1C**). Findings were highly similar, suggesting robustness. The shared disease effect resembled the effects of each condition, with the strongest spatial similarity to SZ and BD, followed by MDD, ADHD, ASD, and OCD (spin-test followed by false discovery rate (FDR) correction, p_spin-FDR_ < 0.001; **Fig. S2**), indicating that the shared effect captured structural alterations from each condition. We furthermore re-evaluated the shared dimension using leave-one-condition-out procedure (see *Methods*), and observed largely consistent results with the shared effect based on all conditions (r > 0.9 p_spin-FDR_ < 0.001; **Fig. S3**), indicating that a single condition with strong meta-analytic profile did not determine the shared disease effect.

**Fig. 1.**
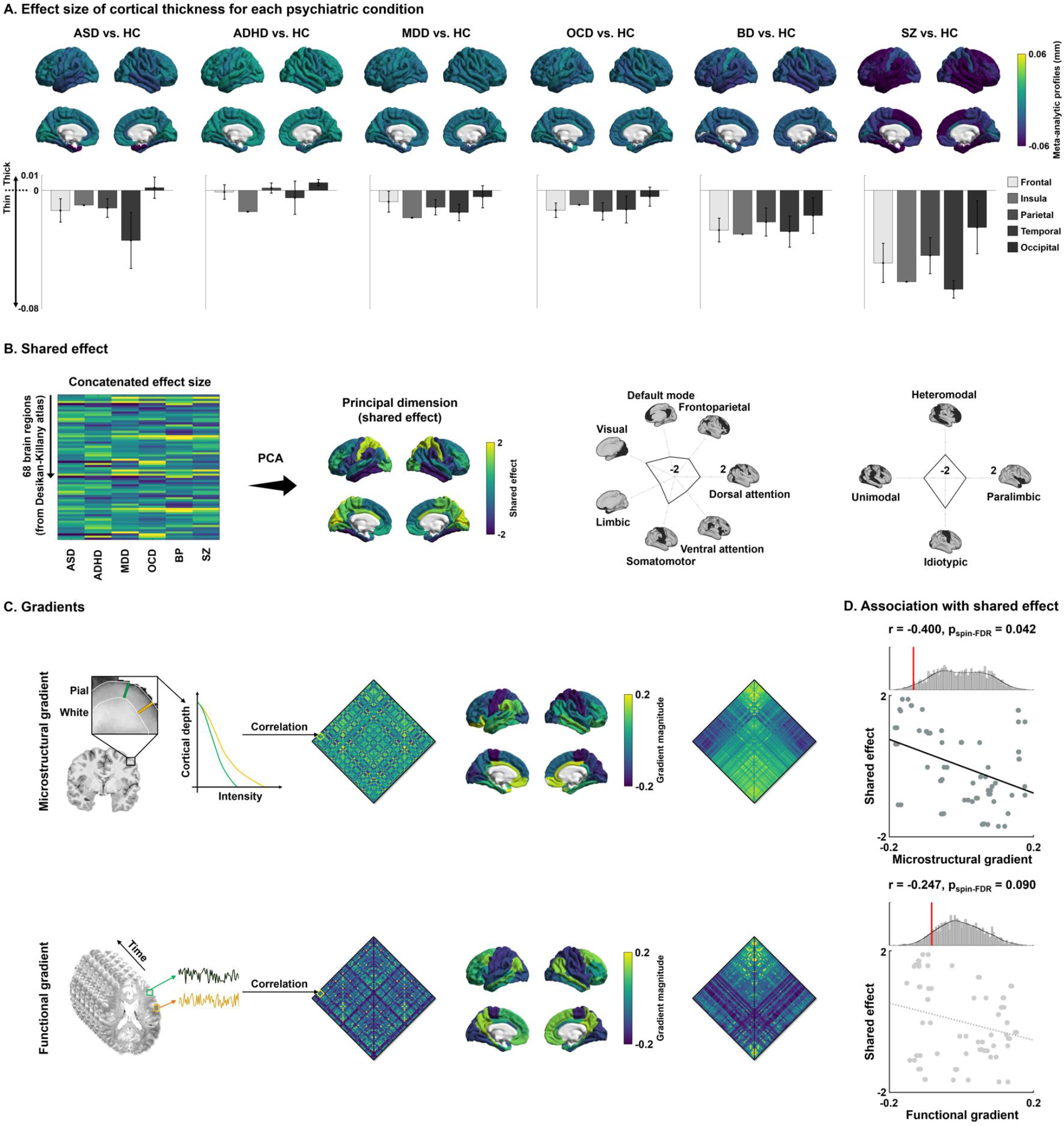
Shared disease effect and associations to connectivity gradients. **(A)** Meta-analytic profiles of cortical thickness differences (unit in mm) in patients with each psychiatric condition relative to matched controls. Positive/negative values indicate increases/decreases in cortical thickness in patients relative to controls. Mean values of the regions involved in the same cortical lobes with SD are reported with bar plots. **(B)** The shared effect was identified through principal component analysis (PCA) applied to the concatenated effect size map. Spider plots stratify the effects according to functional communities [63] and cortical hierarchy levels [40]. **(C)** The microstructural and functional connectivity gradients were generated by applying non-linear dimensionality reduction techniques to the group averaged connectivity matrix (middle left), and each connectivity matrix was reordered (right) according to the first gradients (middle right). **(D)** Spatial correlation of each gradient with the shared effect map are shown in the scatter plots. The distribution of correlation coefficients across 1,000 spin-tests are reported with histograms, and the actual r-values are represented with red bars. *Abbreviations:* ASD, autism spectrum disorder; ADHD, attention deficit hyperactivity disorder; MDD, major depressive disorder; OCD, obsessive-compulsive disorder; BD, bipolar disorder; SZ, schizophrenia; HC, healthy controls.

### Associations with cortical myeloarchitecture and functional connectivity gradients

To assess *in vivo* micro- and macroscopic properties of the shared disease dimension on cortical morphology, we first examined its association with myeloarchitecture and intrinsic functional connectivity gradients [37, 39] (see *Methods*; **Fig. 1C**). The microstructural gradient was derived from inter-regional similarity matrices of intracortical profiles of myelin-sensitive MRI [37], and runs from sensory/motor regions with high laminar differentiation and high intracortical myelin content towards paralimbic cortices with reduced laminar differentiation and low myelin content [37]. The intrinsic functional gradient was derived from resting-state functional MRI connectivity. While it also runs from sensory to transmodal areas, it finds its apex in the heteromodal default mode and frontoparietal networks, and not in paralimbic cortices [39]. Associating the patterns of shared dimension with these two *in vivo* gradients, we observed a negative association with the microstructural gradient (r = -0.400, p_spin-FDR_ = 0.042) and a negative trend with the functional connectivity gradient (r = -0.247, p_spin-FDR_ = 0.090; **Fig. 1C**). In other words, transdiagnostic morphological alterations follow sensory-fugal gradients of cortical organization, in particular the microstructural gradient that differentiates sensory/motor areas with high myelination and distinct lamination from paralimbic areas with low myelin content and reduced laminar differentiation.

### Cytoarchitectonic associations

We furthermore examined associations with cortical cytoarchitecture [36], using a 3D histological reconstruction of a *post mortem* human brain, the BigBrain [60, 64]. We calculated cortex-wide variations in cytoarchitecture using two alternative approaches. First, we obtained intracortical intensity profiles and calculated their statistical moments, *i*.*e*., mean, SD, skewness, and kurtosis (**Fig. 2A-B**). In both classic cytoarchitecture analysis and more recent work, these features have been shown to relate to inter-areal microstructural differentiation [38, 65]. For example, the skewness moment describes spatial transition from areas with low laminar differentiation and negative skewness to those with high laminar differentiation and positive skewness [65–67]. Moreover, we computed externopyramidization [68], describing gradual shift of intensity profiles across cortical layers that has been suggested to differentiate areas on the lower end of the cortical hierarchy from those that are higher up due to hierarchical shifts in laminar projection profiles [69] (**Fig. 2A-B**). Spatial correlations between these features and the principal disease dimension indicated relations to both profile skewness (r = 0.400, p_spin-FDR_ = 0.015) and externopyramidization (r = 0.472, p_spin-FDR_ = 0.015; **Fig. 2C**). In other words, transdiagnostic alteration in cortical morphology was more likely in paralimbic regions with low skewness and low externopyramidization, independently confirming that those areas with low laminar differentiation were more likely to show transdiagnostic cortical alterations.

**Fig. 2.**
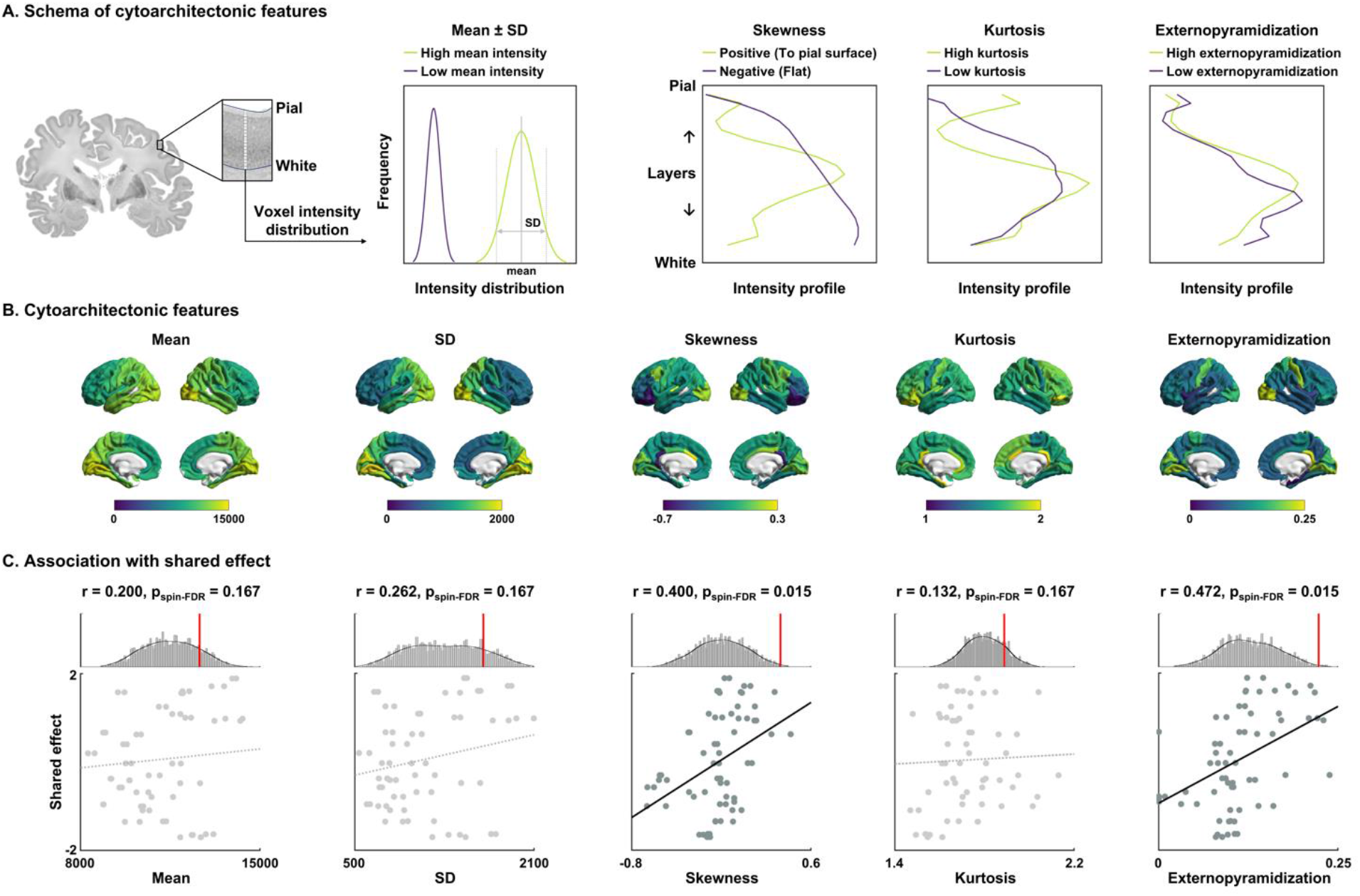
Cytoarchitectonic associations with the shared disease effect. **(A)** Cytoarchitectonic moment features of mean, SD, skewness, and kurtosis, as well as externopyramidization of intracortical intensity profile were calculated from the *post mortem* human brain, and (B) plotted on brain surfaces. **(C)** Spatial correlations between the features and shared effects are shown on scatter plots. The distributions of correlation coefficients across 1,000 spin-tests are reported with histograms, and the actual r-values are represented with red bars. *Abbreviation:* SD, standard deviation.

### Associations with distributions of neurotransmitter systems

Neurotransmitter contextualization leveraged JuSpace [52], a toolbox that disseminates *in vivo* PET/SPECT data sensitive to ten different transmitters/transporters/receptors from independent studies in healthy human adults [53–58] (**Fig. 3A**). Associating the shared dimension with cortex-wide neurotransmitter maps, we observed positive associations with D2 and 5-HT1b receptor densities (D2: r = 0.280, p_spin-FDR_ = 0.035; 5-HT1b: r = 0.349, p_spin-FDR_ = 0.025), and negative correlations with dopamine transporter and 5-HT1a receptor density (DAT: r = -0.240, p_spin-FDR_ = 0.041; 5-HT1a: r = -0.307, p_spin-FDR_ = 0.033; **Fig. 3B**). The results indicate that common cortical abnormality patterns across psychiatric and neurodevelopmental conditions may be reflected by serotonergic and dopaminergic systems. More specifically, higher transdiagnostic cortical atrophy was related to higher 5-HT1a and lower 5-HT1b, as well as higher DAT and lower D2 receptor density.

**Fig. 3.**
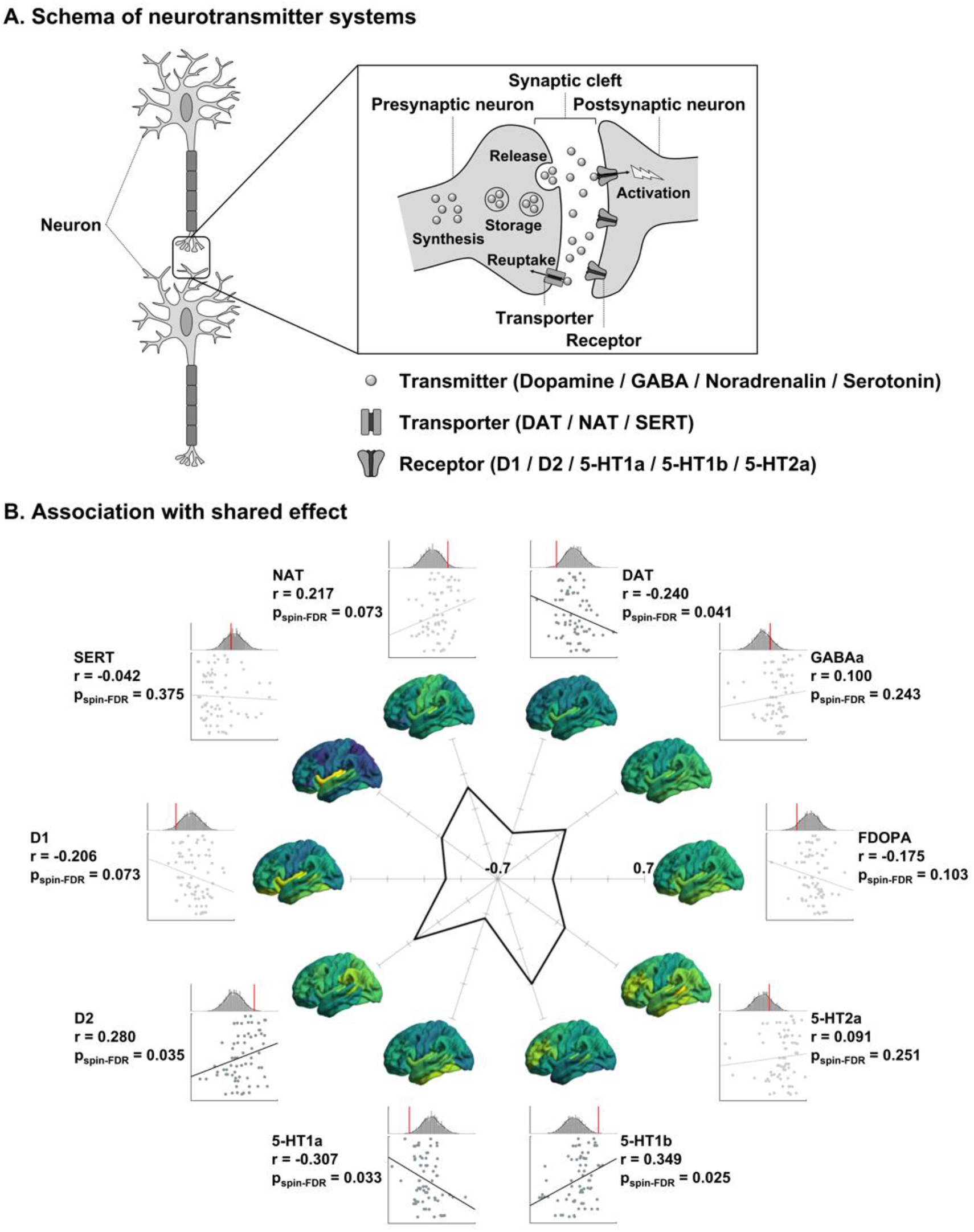
Associations of neurotransmitter systems with shared disease effect. **(A)** Schema of neurotransmitter systems of transmitters, transporters, and receptors. **(B)** Spatial correlations of each neurotransmitter map with shared effect are shown on scatter plots. The distributions of correlation coefficients across 1,000 spin-tests are reported with histograms, and actual r-values are reported with red bars. The spider plot shows correlation coefficients. Cortex-wide spatial maps of the transmitter systems are reported on brain surfaces. *Abbreviations:* FDOPA, 18F fluorodopa; DAT, dopamine transporter; NAT, noradrenaline transporter; SERT, serotonin transporter.

### Prediction of the shared disease effect

As a final analysis, we used supervised machine learning to predict the shared dimension using the above multiscale features. Specifically, we leveraged least absolute shrinkage and selection operator (LASSO) regression [70] with five-fold nested cross-validation [71–74] to predict the cross-condition effect using concatenated multiscale features (see *Methods*; **Fig. 4A**). Repeating the analysis for 100 times with different training and test dataset subsplits, we could reliably predict the spatial pattern of the shared disease dimension (mean ± SD r = 0.518 ± 0.044, mean absolute error (MAE) = 0.828 ± 0.039, p_perm_ < 0.001; **Fig. 4B**). Cytoarchitectural skewness and externopyramidization, followed by D2 and 5-HT1b receptors, as well as the microstructural gradient were frequently selected across cross-validations and repetitions (**Fig. 4A**). When considering each psychiatric condition separately, we could find significant prediction performances, but the features selected diverge across conditions (**Fig. S4**).

**Fig. 4.**
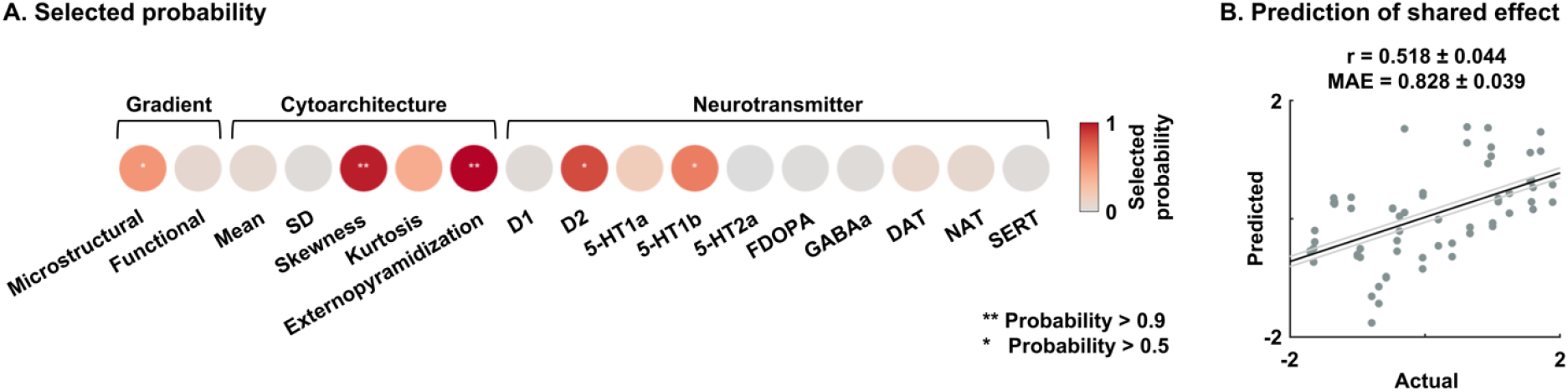
Association between the shared disease effect and multiscale features using machine learning. **(A)** Probability of the selected features across five-fold nested cross-validations and 100 repetitions for predicting the shared disease effect. The frequently selected features are reported with asterisks. **(B)** Linear correlation between actual and predicted values of the effects is shown on a scatter plot. Black line indicates mean correlation and gray lines represent the 95% confidence interval for 100 iterations with different training/test datasets. *Abbreviations:* SD, standard deviation; FDOPA, 18F fluorodopa; DAT, dopamine transporter; NAT, noradrenaline transporter; SERT, serotonin transporter; MAE, mean absolute error.

## Discussion

The current work determined cortex-wide variations in susceptibility to morphological alterations across six major psychiatric conditions (*i*.*e*., ASD, ADHD, MDD, OCD, BD, and SZ), and cross-referenced these spatial patterns against multiscale cortical organization. Specifically, studying data aggregated by several multi-site ENIGMA consortia on the above indications [26–32], we identified a shared morphological dimension that followed a sensory-fugal pattern of increasing susceptibility to morphological alterations in paralimbic regions. Moreover, we cross-referenced these findings against neural axes previously described by (i) *in vivo* MRI measures sensitive to cortical myeloarchitecture and intrinsic functional connectivity [37, 39], (ii) *post mortem* histological measures sensitive to cytoarchitecture, in particular laminar differentiation [36, 38, 60, 65], and (iii) *in vivo* PET/SPECT derived measures of cortical neurotransmitter systems [52–58]. Our findings revealed that the transdiagnostic dimension of morphological anomalies closely aligned with microstructural gradients differentiating sensory/motor from paralimbic areas on the basis of cortical cyto- and myeloarchitecture, together as well as the variable distribution of serotonin and dopamine neurotransmitter systems. By offering new insights into multiscale neural features that align with cortical structural compromise across several psychiatric conditions, our work outlines micro- and macroscale determinants of cerebral vulnerability to the effects of common mental illnesses.

Complementing earlier case-control MRI studies performed separately in common neuropsychiatric conditions [17, 26–31, 75], an emerging literature of both primary observational studies [22–24] as well as meta-analyses [32, 33] has increasingly investigated sets of disorders to explore transdiagnostic effects on brain structure. Recently, these studies were complemented by dimensional data decomposition approaches of cortical morphological data, for example a recent factor analysis [33] and principal component analyses [32]. We extended these prior studies by calculating mean effect size of the previously published condition-specific effects [32], as well as applying principal component analysis to the effect sizes and those sourced from the ENIGMA toolbox [61], confirming robust patterns. Specifically, the first shared dimension of cortical morphological alterations described a gradual axis running from sensory-motor regions at one end towards transmodal and most specifically paralimbic areas on the other end. In a prior study, the shared dimension was cross-referenced against gene expression information from the Allen Brain Institute [76–78], a comprehensive microarray-derived transcriptomics dataset based on four *post mortem* brains in the left and two in both left and right hemispheres. Using this resource, the authors found that transdiagnostic effects were highest in brain regions expressing genes for pyramidal *cornu Ammonis* 1 (CA1) cells, a finding that may already point towards a potentially increased susceptibility of limbic allocortices to transdiagnostic effects on brain morphology [32]. Here, we extended these findings by contextualizing the shared disease effect across multiple scales of neural organization, including cortex-wide variations in myeloarchitecture, cytoarchitecture, intrinsic functional connectivity, as well as neurotransmitter distributions.

The *in vivo* microstructural cortical gradient was defined using a recently-introduced procedure [37], which identified axes of cortico-cortical differentiation based on the similarity of myelin-sensitive MRI profiles sampled across cortical depths. In healthy adults and adolescents [37, 65], this approach has revealed a robust sensory-fugal cortical gradient running from sensory/motor areas with marked laminar differentiation and high myelin content towards paralimbic cortices with low overall myelination and rather agranular cortical profiles. By showing an association between the shared dimension and this microstructural gradient, we confirm an overall heightened susceptibility of paralimbic cortices to disease-related cortical thickness changes. Several architectonic features of the paralimbic cortices may underscore their increased susceptibility to disease-related effects. On the one hand, these regions have an architecture that may permit an increased potential for brain plasticity. This includes an overall reduced neuronal density in paralimbic regions compared to eulaminate cortices that may be more permissive for dendritic arborization and synaptogenesis [34]. Paralimbic areas also express several developmental markers into adulthood that cease to be expressed in other areas after ontogeny, such as growth associated protein GAP-43 [79]. On the other hand, limbic areas are known to have a relatively late myelination compared to sensory/motor areas and lower overall myelin content in adults. The role of intracortical myelination in plasticity is likely to be complex, but several streams of evidence point to a role of myelin acting as a buffer against plasticity. In addition to acting as an insulator for electrical transmission, myelin associated growth inhibitors limit activity and experience-induced axon sprouting, with downstream effects of synaptic plasticity [80]. The reduced myelin content, together with increased complexity of dendritic arborization in transmodal and paralimbic regions may render cortical microstructure in these regions more susceptible to pathological alterations, which would echo observations in other neurological conditions. For example, the core pathological substrates of drug-resistant temporal epilepsy is thought to be localized in limbic/paralimbic regions [81–83], and prior work has suggested rather specific changes in myelin and microstructural proxies in these areas [84, 85]. Similar findings have been observed in degenerative conditions such as Alzheimer’s disease [44, 86, 87] as well as depression [88] and autism [89, 90], where pathology spreads from disease epicenters in paralimbic allocortices to invade more widespread cortical/subcortical networks. These findings collectively show that cellular and molecular features of paralimbic cortices and their cortico-cortical pathways promote brain plasticity as well as higher metabolic activity, and are thus likely more vulnerable to both developmental as well as acquired disruptions than other regions, supporting the hypothesis that their cortical type predisposes to a heightened vulnerability for an impact of neuropsychiatric conditions on alterations in brain morphology [34].

Studying the *post mortem* 3D BigBrain [60], we obtained supporting confirmation for the above association between cortical microstructure and disease related susceptibility. In particular, we discovered similarly marked associations between the shared disease dimension and laminar profile skewness as well as externopyramidization, both features assessing depth-dependent shifts in the distribution in cell densities [38, 68]. In prior work, we reported that the profile skewness feature discriminates unimodal granular cortices from agranular/dysgranular paralimbic regions at a cortex-wide level [65], and also helped to delineate the iso-to-allocortical axis in the mesiotemporal lobe system [66]. Studying typical adolescent development, changes in profile skewness of myelin-sensitive MRI contrasts have furthermore been reported to spatially co-localize with expression patterns of genes enriched in oligodendrocytes [65]. As a complementary feature of laminar organization, externopyramidization classically contextualizes the ratio of neuronal densities between supragranular and infragranular cortical layers. It increases when the cortex is cytoarchitectonically more differentiated, which happens in primary areas with a marked layer 4 [68]. Thus, the association of these cortical depth-dependent cytoarchitectural features with the shared disease effect confirms the *in vivo* findings with ultrahigh resolution cytoarchitecture data suggesting that paralimbic areas, sensitive to transdiagnostic cortical alterations, are less laminarly differentiated. Furthermore, prior cellular and transcriptomic studies indicate regional susceptibility of synaptic elements as well as mutated genes in schizophrenia [91, 92] and bipolar disorder [93]. Indeed, major depression may cause atrophy of neurons in limbic regions [94], pointing histopathological susceptibility of paralimbic areas in psychiatric conditions.

We also observed a marginal association between the transdiagnostic effect on brain structure and the principal functional connectivity gradient, but findings were overall weaker than for the above *in vivo* and *post mortem* derived microstructural gradients. Motifs of macroscale intrinsic functional connectivity also show an overall sensory-fugal pattern [40, 95–97], but the associated gradients generally run from sensory/motor towards more heteromodal association cortices such as the default mode and frontoparietal networks, and not the paralimbic regions. These findings may indirectly support the conclusion that transdiagnostic disease effects on brain morphology may more closely align with spatial trends in microstructure rather than with macroscale functional differentiation. As brain organization show functional heterogeneity and multiplicity, investigation of associations between the transdiagnostic effects and multiple functional gradients is required for further studies. Notably, however, the cortical morphological data from the ENIGMA dataset were only available in the Desikan-Killany parcellation [62], a relatively macroscopic scheme mainly based on sulco-gyral features. In addition to not offering a high granularity on cortical arealization, the reliance on folding alone may only provide rather limited sensitivity to contextualize our findings with respect to functional topographies. It would thus be relevant to re-evaluate functional gradient association based on functionally-defined parcellations [98, 99] or at a vertex-level.

In addition to our findings showing overall associations between the transdiagnostic effect and sensory-fugal microstructural gradients, we observed associations to the spatial distribution of different neurotransmitter systems derived from *in vivo* neuroimaging. Notably, associations were seen both to serotonin (5-HT1a and 5-HT1b) and dopamine receptors and transporters (DAT/D1 and D2), two important markers of mental health and targets for pharmacological treatments [100–108]. In both cases (*i*.*e*. 5-HT1a vs 5-HT1b, DAT/D1 vs D2), associations to the disease effect were of opposite polarity, confirming prior work in rodents [109–113] and humans [114–117]. Associations with *in vivo* neurotransmitter topographies provide a novel way of indirectly assessing the relationship between shared abnormalities on cortical morphology and neurotransmitter systems so that we can understand putative mechanisms of shared morphological abnormalities, extending prior work from rodents and humans. As an integrative analysis, we opted for supervised machine learning to predict the shared disease effect. This analysis revealed that not a single feature, but rather combinations of both microstructure and dopamine/serotonin transmitter systems have highest utility in predicting the spatial pattern of the transdiagnostic morphological dimensions. Overall, our findings add new evidence for a principal organizational dimension that differentiates sensory-motor networks from transmodal cortices in typical human brain organization [37, 39, 118], and furthermore describes the main axis of cortex-wide susceptibility to transdiagnostic effects of common mental health conditions. Altogether, the observed associations between multiscale neural mechanisms and transdiagnostic anomalies of cortical morphology provide a potentially integrative framework for understanding neuropathology in psychiatry and the development of treatment that cut across traditional disease boundaries.

## Methods

### Study dataset

#### (a) ENIGMA data

We analyzed T1-weighted data from people with a diagnosis of (n = 12,876) ASD (n = 1,821), ADHD (n = 1,815), MDD (n = 2,695), OCD (n = 2,274), BD (n = 1,555), and SZ (n = 2,716) and site matched healthy controls (n = 15,670) from 145 independent cohorts participating in prior ENIGMA consortium studies [26–31]. Demographic information is summarized in **Table S1** and available in a recent cross-condition study [32]. Data from each center were processed using the standard ENIGMA workflow (http://enigma.ini.usc.edu/protocols/imaging-protocols/). Processing was conducted using FreeSurfer [119–121] that involves magnetic field inhomogeneity correction, non-brain tissue removal, intensity normalization, and tissue segmentation. Estimated white and pial surfaces were inflated to spheres and registered to the *fsaverage* template. Based on the Desikan-Killiany atlas [62], cortical thickness was measured for 68 gray matter brain regions. For each psychiatric condition, the ENIGMA groups performed multiple linear regression analyses to fit cortical thickness measures with age, age squared, sex, and site information. The meta-analytic profiles of between-group differences between patients and controls were estimated via an inverse variance-weighted random-effects model, which can be obtained from the previous study [32] (**Fig. 1A**). If the studies provided multiple effect sizes across children/adolescents/adults, only the effects from the adult sample were used, in order to match the age range across conditions. The positive/negative effects indicate increases/decreases in cortical thickness in patients relative to controls. Individual cohort investigators obtained approval from local institutional ethics boards, and informed consent was obtained from study participants or their guardians.

#### (b) HCP data

To generate microstructural and functional connectivity gradients, we also studied 207 unrelated healthy young adults (60% females, mean age ± SD = 28.73 ± 3.73 years) from the HCP dataset [59]. In the HCP, multimodal imaging data comprising T1- and T2-weighted as well as rs-fMRI were acquired on a Siemens Skyra 3T at Washington University. The cohort selection is identical to our prior work [61, 122]. T1-weighted images were acquired using a magnetization-prepared rapid gradient echo (MPRAGE) sequence (repetition time (TR) = 2,400 ms; echo time (TE) = 2.14 ms; inversion time (TI) = 1,000 ms; flip angle = 8°; field of view (FOV) = 224 × 224 mm^2^; voxel size = 0.7 mm isotropic; 256 slices). T2-weighted data were obtained using a T2-SPACE sequence, with the same acquisition parameters as for the T1-weighted data except for TR (3,200 ms), TE (565 ms), and flip angle (variable). The rs-fMRI data were collected using a gradient-echo echo-planar imaging sequence (TR = 720 ms; TE = 33.1 ms; flip angle = 52°; FOV = 208 × 180 mm^2^; voxel size = 2 mm isotropic; number of slices = 72; and 1,200 volumes per time series), where participants were instructed to keep their eyes open looking at a fixation cross during the scan. Two sessions (left-to-right and right-to-left phase-encoded directions) of rs-fMRI data were acquired, providing up to four time series per participant.

Images underwent minimal preprocessing pipelines using FSL, FreeSurfer, and Workbench as follows [123–125]:

#### i) T1- and T2-weighted data

Data were corrected for gradient nonlinearity and b0 distortions, and then T1- and T2-weighted data were co-registered using a rigid-body transformation. Bias field was adjusted based on the inverse intensities from the T1- and T2-weighting. The white and pial surfaces were generated [119–121], and the mid-thickness surface was generated by averaging them. The mid-thickness surface was inflated and the spherical surface was registered to the Conte69 template with 164k vertices [126] using MSMAll [99] and downsampled to a 32k vertex mesh.

#### ii) Microstructure data

Myelin-sensitive proxy was estimated based on the ratio of the T1- and T2-weighted contrast [127, 128]. We generated 14 equivolumetric surfaces within the cortex and sampled T1w/T2w intensity along these surfaces [37]. A microstructural similarity matrix was constructed by calculating linear correlation of cortical depth-dependent T1w/T2w intensity profiles between different cortical regions based on the Desikan-Killiany atlas [62], controlling for the average whole-cortex intensity profile [37]. The matrix was thresholded at zero and log-transformed [37]. A group matrix was constructed by averaging matrices across participants.

#### iii) rs-fMRI data

Data were corrected for distortions and head motion, and registered to the T1-weighted data and subsequently to MNI152 standard space. Magnetic field bias correction, skull removal, and intensity normalization were performed. Noise components attributed to head movement, white matter, cardiac pulsation, arterial, and large vein related contributions were removed using FMRIB’s ICA-based X-noiseifier (ICA-FIX) [129]. Preprocessed time series were mapped to the standard ‘grayordinate’ space using a cortical ribbon-constrained volume-to-surface mapping algorithm. The total mean of the time series of each left-to-right/right-to-left phase-encoded data was subtracted to adjust the discontinuity between the two datasets and then concatenated to form a single time series. A functional connectivity matrix was constructed by calculating the linear time series correlations between Desikan-Killiany parcels [62], followed by Fisher’s r-to-z transformation [130]. Individual connectivity matrices were averaged to construct a group level connectome.

### Shared effects of cortical thickness differences across conditions

To assess transdiagnostic effects of cortical thickness differences in patients relative to controls, we applied principal component analysis to the concatenated effect size maps across six conditions [131] (**Fig. 1B** and **Fig. S1A**). The first principal dimension was determined as the shared disease effect. We summarized the effects according to seven intrinsic functional communities [63], as well as four cortical hierarchical levels [40]. We additionally calculated mean effect size across the conditions to intuitively interpret shared disease effect (**Fig. S1B**), and also estimated principal dimension based on the data sourced from the ENIGMA toolbox (*i*.*e*., Cohen’s d; **Fig. S1C**). We compared the shared dimension and the effect size of each condition via linear correlations to assess the degree of contribution of each condition (**Fig. S2**). The significance of the correlation was determined using 1,000 non-parametric spin-tests, to account for spatial autocorrelation [132], and corrected for multiple comparisons using a FDR procedure [133]. To assess robustness, we performed leave-one-condition-out cross-validation. Specifically, we estimated the shared dimension using five conditions by excepting for a single condition, and assessed similarity with the shared disease effect estimated based on the whole six conditions (**Fig. S3**). We calculated significance of the correlation using 1,000 spin-tests, and multiple comparisons were corrected using FDR [132, 133].

### Associations to microstructural and functional connectivity gradients

We evaluated the underlying connectome organizations of the shared disease effects. Based on T1w/T2w and rs-fMRI data obtained from the HCP database [59], we estimated microstructural and functional gradients, the low dimensional representation of connectome organizations explaining spatial variation in the connectome data [37, 39], using BrainSpace (https://github.com/MICA-MNI/BrainSpace) [97] (**Fig. 1C**). An affinity matrix was constructed with a normalized angle kernel from the group averaged connectivity matrix with the top 10% entries for each parcel. The connectome gradients were estimated using diffusion map embedding [134], which is robust to noise and computationally efficient compared to other non-linear manifold learning techniques [71, 135]. It is controlled by two parameters α and t, where α controls the influence of the density of sampling points on the manifold (α = 0, maximal influence; α = 1, no influence) and t scales eigenvalues of the diffusion operator. The parameters were set as α = 0.5 and t = 0 to retain the global relations between data points in the embedded space, following prior applications [37, 39, 47, 97, 136]. We associated the shared effect with these gradients using linear correlation (**Fig. 1D**), where the significance was assessed using 1,000 spin-tests followed by FDR [132, 133].

### Cytoarchitectonic associations with shared disease effects

We aimed to associate the shared dimensions with histology-driven cytoarchitectonic features derived from BigBrain surfaces with 62 cortical areas (https://bigbrain.loris.ca/main.php) [60]. Specifically, BigBrain is a ultra-high resolution, 3D volumetric reconstruction of a *post mortem* Merker-stained and sliced human brain from a 65-year-old male, with specialized pial and white matter surface reconstructions [60]. The *post mortem* brain was paraffin-embedded, coronally sliced into 7400 20-μm sections, silver-stained for cell bodies [137], and digitized. A 3D reconstruction was implemented with a successive coarse-to-fine hierarchical procedure, resulting in a full brain volume. Among 68 regions defined by the Desikan-Killiany atlas [62], three regions per hemisphere, including banks of superior temporal sulcus, frontal pole, and temporal pole, were excluded as the BigBrain did not provide data for these regions. We generated 18 equivolumetric cortical surfaces within the cortex (https://github.com/caseypaquola/BigBrainWarp) and sampled the intensity values along these surfaces. Based on the intensity values, we calculated four moment features, including mean, SD, skewness, and kurtosis, as well as externopyramidization (**Fig. 2A-B**). The mean and SD represent the overall intensity distribution of cytoarchitecture across layers, skewness indicates shifts in intensity values towards supragranular layers (*i*.*e*., positive skewness) or flat distribution (*i*.*e*., negative skewness), and kurtosis identifies whether the tails of the intensity distribution contain extreme values. Externopyramidization reflects gradual shifts of intensity values from infragranular to supragranular layers defined as follows [69]:

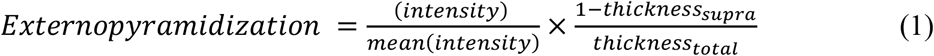

To assess associations with shared disease effects, we calculated linear correlations between cytoarchitectonic features and shared effects (**Fig. 2C**). The significance of the correlations was assessed using 1,000 spin-tests followed by FDR across different cytoarchitectonic features [132, 133].

### Associations between transmitter systems and shared effects

To provide underlying molecular properties of the shared effects in neuroanatomical disruptions across different psychiatric conditions, we associated the shared dimensions with ten different neurotransmitter maps of healthy controls provided by prior independent PET/SPECT studies [53– 58], which contain neurotransmitters of FDOPA, GABAa, transporters of DAT, NAT, SERT, and receptors of D1, D2, 5-HT1a, 5-HT1b, and 5-HT2a (https://github.com/juryxy/JuSpace) [52] (**Fig. 3A**). All PET maps were linearly rescaled to have intensity values between 0 and 100 [52]. After mapping the neurotransmitter maps onto the Desikan-Killiany atlas [62], we calculated linear correlations between the shared effects and each neurotransmitter map (**Fig. 3B**), and assessed the significance using 1,000 spin tests followed by FDR to adjust for multiple comparisons across ten different maps [132, 133].

### Prediction of shared effects using multiscale features

We associated multiscale features and shared effects using supervised machine learning to incorporate our findings (**Fig. 4**). Specifically, we aimed to predict the shared disease effects using concatenated multiscale features of microstructural and functional gradients, cytoarchitectonic (*i*.*e*., mean, SD, skewness, kurtosis, externopyramidization), and transmitter maps (*i*.*e*., D1, D2, 5-HT1a, 5-HT1b, 5-HT2a, FDOPA, GABAa, DAT, NAT, SERT). We used five-fold nested cross-validation [72–74] with LASSO regression [70]. Nested cross-validation split the dataset into training (4/5) and test (1/5) partitions, and each training partition was further split into inner training and testing folds using another five-fold cross-validation. The model with the best performance (lowest MAE) across the inner folds was applied to the test partition of the outer fold. Among the multiscale features, we selected performant features using LASSO regularization, and the effect size was predicted using linear regression with the selected features. The procedure was repeated 100 times with different training and test partitions. Prediction accuracy was evaluated with linear correlations between the actual and predicted effect size and the MAE, with their 95% confidence interval. Permutation-based correlations across 1,000 tests were conducted by randomly shuffling cortical regions to verify whether the prediction performance exceeded chance levels. We also performed the prediction analysis using the effect size of each condition (**Fig. S4**).

## Acknowledgments

The authors would like to express their gratitude to the open science initiatives that made this work possible: (i) The ENIGMA-Epilepsy consortium and in particular the ASD, ADHD, MDD, OCD, BD, and SZ working groups (http://enigma.ini.usc.edu/ongoing/) and (ii) The Human Connectome Project (Principal Investigators: David Van Essen and Kamil Ugurbil; 1U54MH091657) funded by the 16 NIH Institutes and Centers that support the NIH Blueprint for Neuroscience Research; and by the McDonnell Center for Systems Neuroscience at Washington University.

## Funding

Bo-yong Park was funded by the National Research Foundation of Korea (NRF-2021R1F1A1052303), Institute for Information and Communications Technology Planning and Evaluation (IITP) funded by the Korea Government (MSIT) (2020-0-01389, Artificial Intelligence Convergence Research Center, Inha University; 2021-0-02068, Artificial Intelligence Innovation Hub), and Institute for Basic Science (IBS-R015-D1). Valeria Kebets was funded by the Quebec Autism Research Training Fellowship of the Transforming Autism Care Consortium (TACC). Sara Larivière was funded by the Canadian Institutes of Health Research (CIHR). Meike D. Hettwer was funded by the Max Planck society and the German Federal Ministry of Education and Research (BMBF). Martine Hoogman is supported by a personal Veni grant from the Netherlands Organization for Scientific Research (NWO, grant number 91619115). Lianne Schmaal was funded by NHMRC Career Development Fellowship (1140764), National Institute of Mental Health of the National Institutes of Health R01MH117601. Ole A. Andreassen was funded by Research Council of Norway (#223273, #283798), KG Jebsen Stiftelsen. Dan J Stein is funded by the SA MRC. Dr. Matthias Kirschner acknowledges support from the National Bank Fellowship (McGill University) and the Swiss National Foundation (P2SKP3_178175). Jessica Turner was funded by National Institutes of Health (NIH 5R01MH121246). Paul Thompson and Sophia Thomopoulos are funded in part by the U.S. National Institutes of Health, under grants R01MH116147, R01MH111671, R01AG058854, RF1MH123163 and U54 EB020403. Boris C. Bernhardt acknowledges research support from the National Science and Engineering Research Council of Canada (NSERC Discovery-1304413), the CIHR (FDN-154298, PJT), SickKids Foundation (NI17-039), Azrieli Center for Autism Research (ACAR-TACC), BrainCanada, Fonds de la Recherche du Québec – Santé (FRQ-S), and the Tier-2 Canada Research Chairs program. Alan Evans and Boris C. Bernhardt were furthermore supported by the Helmholtz International BigBrain Analytics and Learning Laboratory (HiBALL).

## Conflict of interest

Ole A. Andreassen received speaker’s honorarium from Lundbeck and Sunovion, Consultant to HealthLytix. Paul M. Thompson received grant support from Biogen, Inc., and consulting payments from Kairos Venture Capital, for work unrelated to the current manuscript. Other authors declare no conflicts of interest.

## Supporting Information

**Table S1.** Demographic information of studied participants.

| Condition | Number* | Mean (SD; range) age (years) | Sex (male:female) |
| --- | --- | --- | --- |
| ASD/controls | 1821/1823 | 15.6 (6.7; 2–64) | 2941:703 (19% female) |
| ADHD/controls | 1815/1602 | 21.1 (5.4; 4–74) | 2244:1172 (34% female) |
| MDD/controls | 2695/3627 | 40.9 (10.9; 8–89) | 2665:3657 (58% female) |
| OCD/controls | 2274/2013 | 27.2 (8.0; 5–65) | 2166:2121 (49% female) |
| BD/controls | 1555/3423 | 35.1 (12.0; 8–86) | 2142:2836 (57% female) |
| SZ/controls | 2716/3272 | 33.9 (10.7; 7–87) | 3479:2509 (42% female) |

Detailed information available in eTable 1 and eTable 2 of Patel et al., 2021. *Abbreviations:* SD, standard deviation; ASD, autism spectrum disorder; ADHD, attention deficit hyperactivity disorder; MDD, major depressive disorder; OCD, obsessive-compulsive disorder; BD, bipolar disorder; SZ, schizophrenia.

**Fig. S1.**
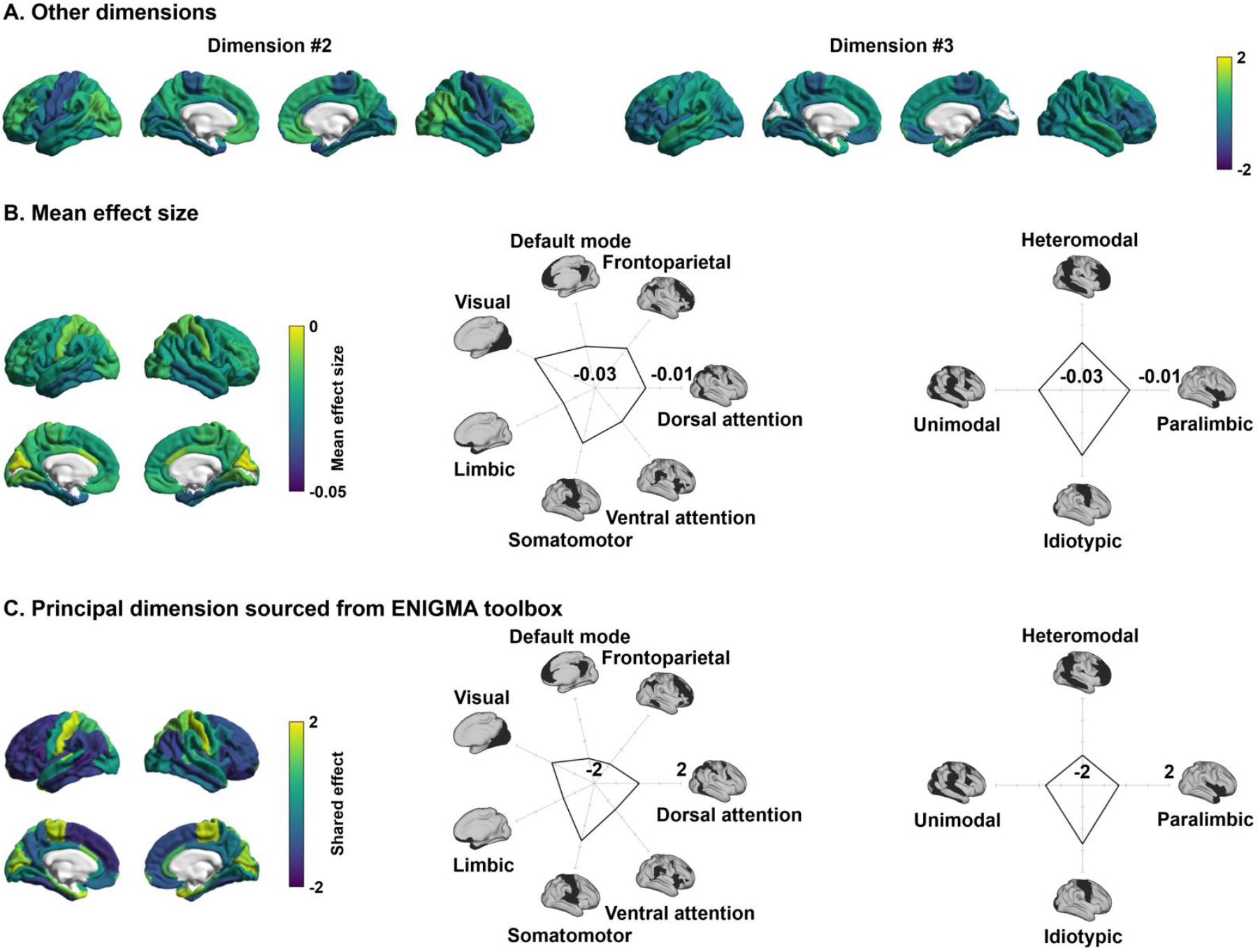
Shared disease effect. **(A)** The second and third dimensions of shared disease effect. **(B)** Mean effect size of cortical thickness alterations across conditions. **(C)** Principal dimension based on the effect size maps (Cohen’s d) sourced from the ENIGMA toolbox. The effects were stratified according to functional communities and cortical hierarchy levels.

**Fig. S2.**
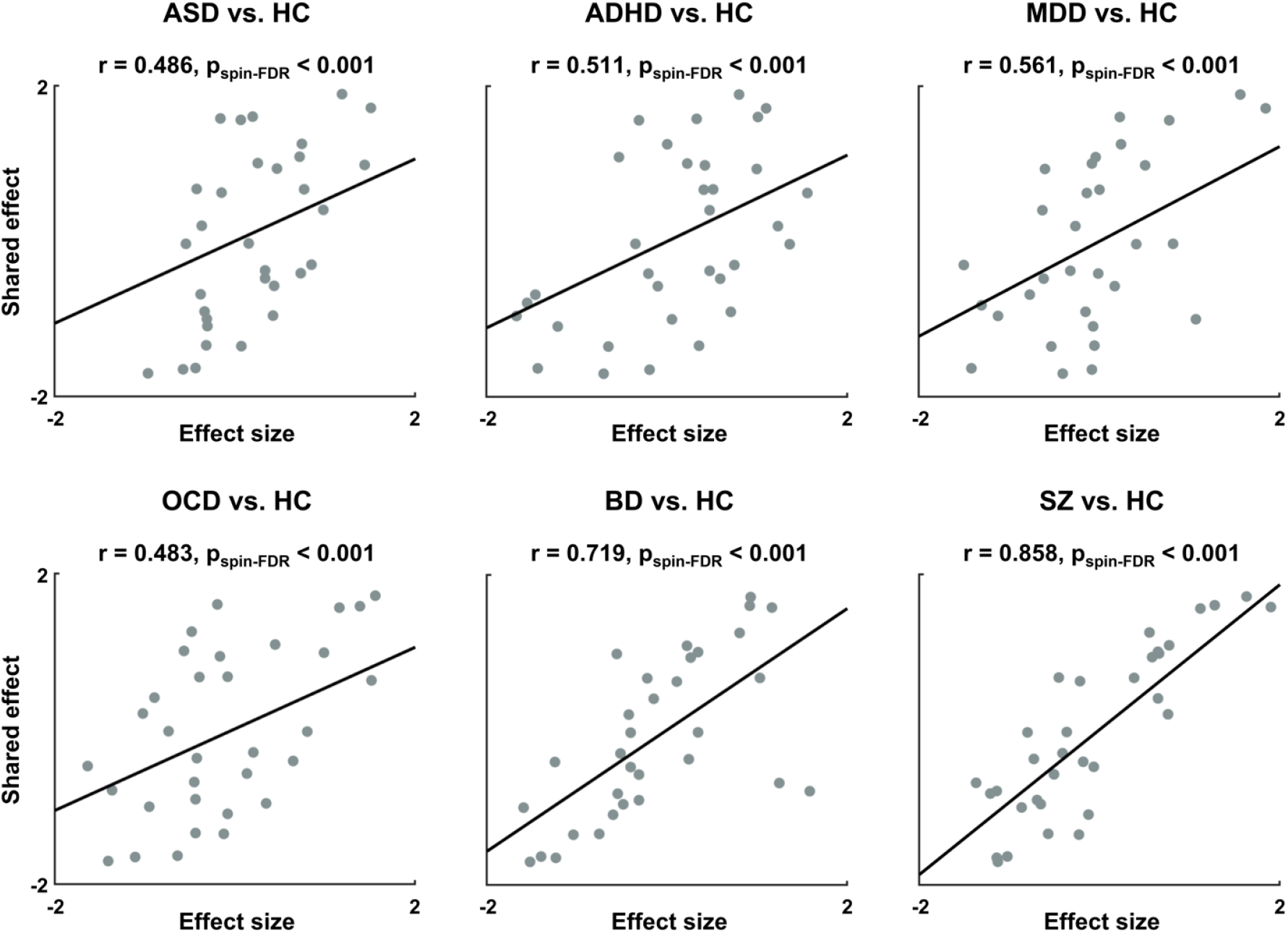
Linear correlations between the shared disease effect and cortical thickness alterations of each condition. *Abbreviations:* ASD, autism spectrum disorder; ADHD, attention deficit hyperactivity disorder; MDD, major depressive disorder; OCD, obsessive-compulsive disorder; BD, bipolar disorder; SZ, schizophrenia; HC, healthy controls.

**Fig. S3.**
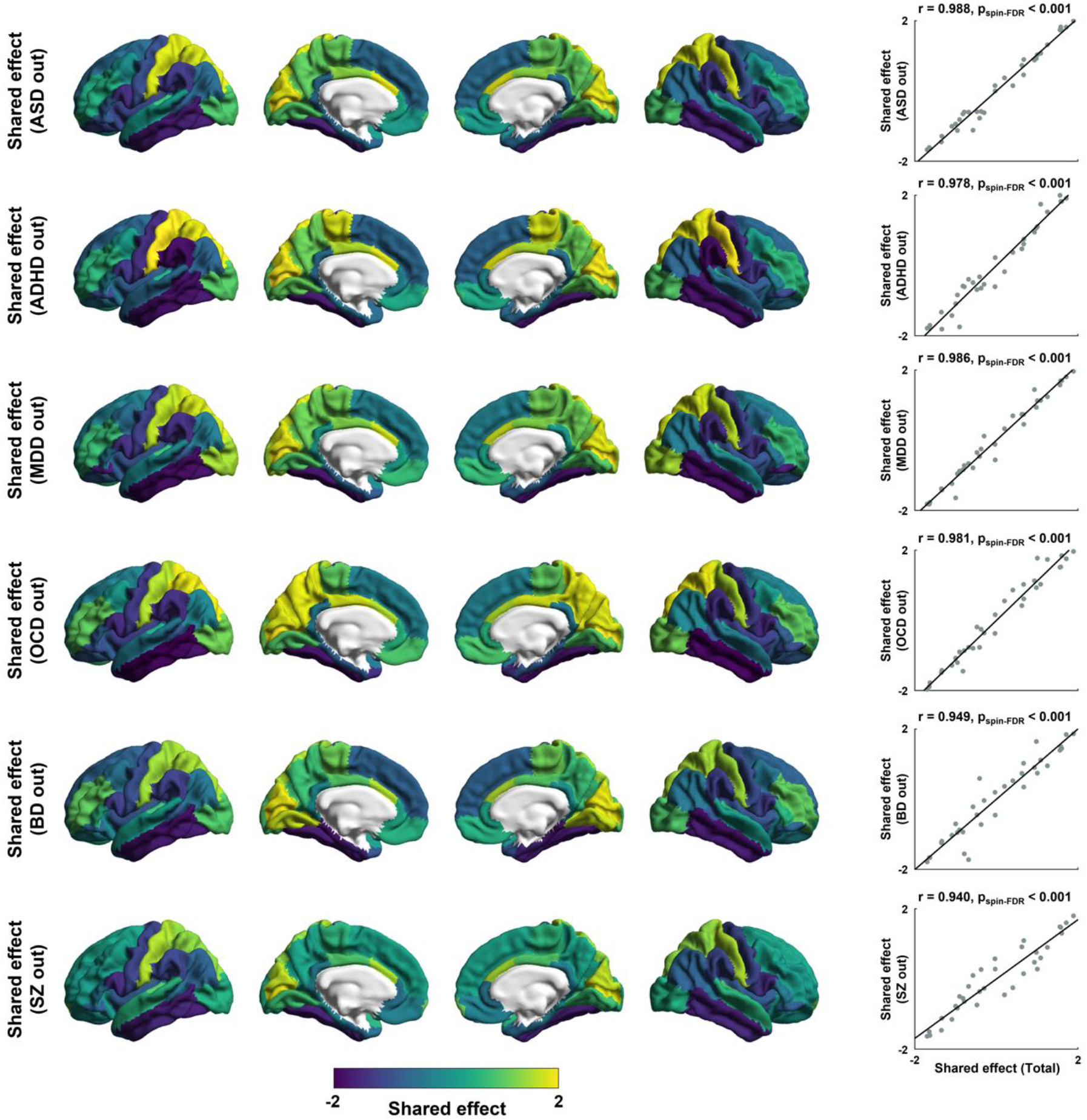
Shared disease effects with leave-one-condition-out cross-validation. The shared dimensions estimated based on all conditions without a single condition are reported on brain surfaces. Linear correlations between the shared effect based on all conditions (see *Fig. 1B*) and that based on five conditions are shown in the scatter plots. *Abbreviations:* ASD, autism spectrum disorder; ADHD, attention deficit hyperactivity disorder; MDD, major depressive disorder; OCD, obsessive-compulsive disorder; BD, bipolar disorder; SZ, schizophrenia; HC, healthy controls.

**Fig. S4.**
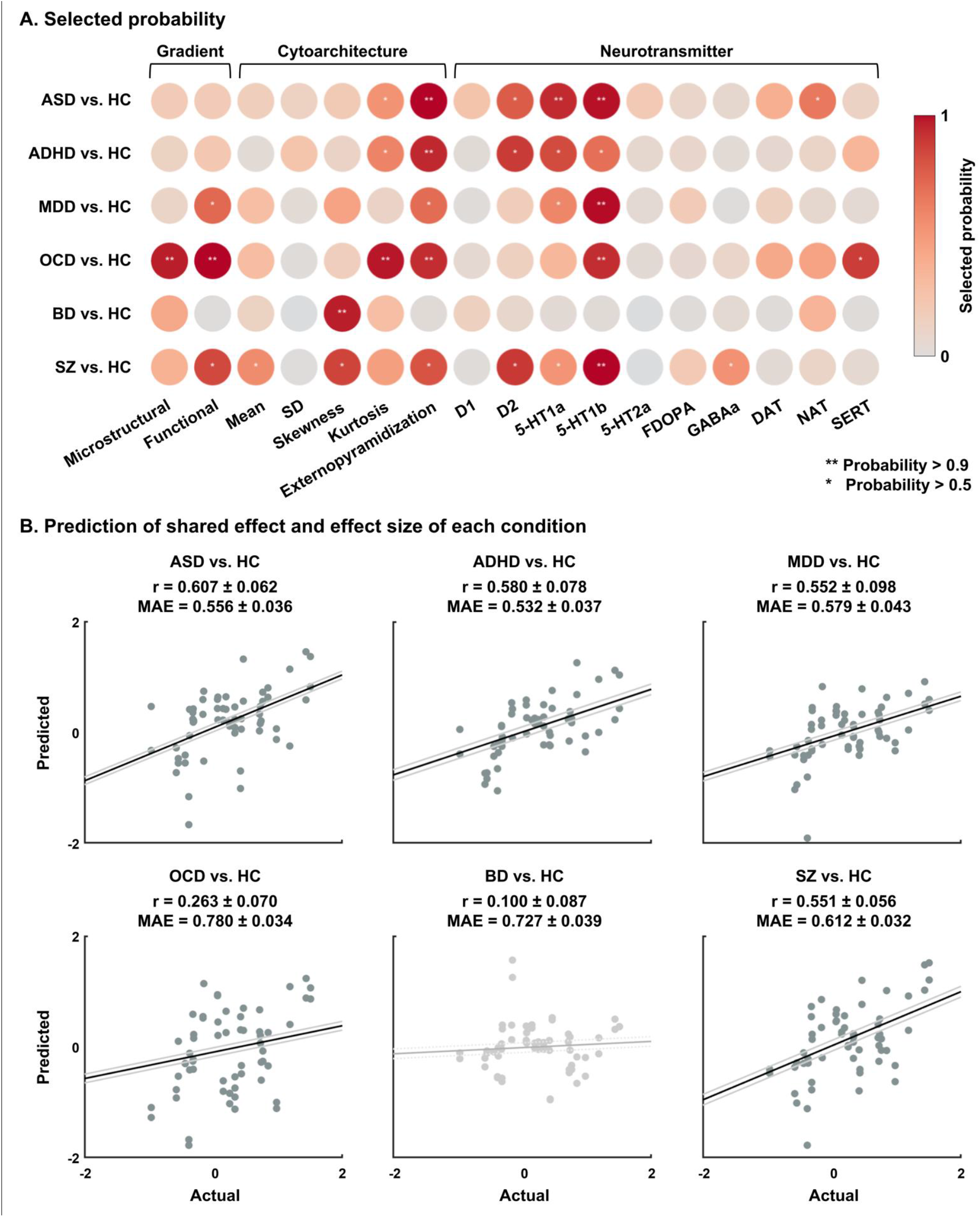
Association between the effect size of each psychiatric condition and multiscale features. **(A)** Probability of the selected features for each psychiatric condition. **(B)** Linear correlations between actual and predicted values of the effects are shown using scatter plots. For details, see *Fig. 4. Abbreviations:* SD, standard deviation; FDOPA, 18F fluorodopa; DAT, dopamine transporter; NAT, noradrenaline transporter; SERT, serotonin transporter; MAE, mean absolute error.

